# Menopause has not evolved as a general trait in mammals: A response to ‘Do mammals have menopause?’

**DOI:** 10.1101/2024.02.29.582687

**Authors:** Simon N. Chapman, Samuel Ellis, Mirkka Lahdenperä, Darren P. Croft, Virpi Lummaa

## Abstract

Reproductive senescence is widespread across mammals, but only a small number of species have physiological reproductive cessation and an extended post-reproductive lifespan. A recent commentary in Cell by Winkler & Goncalves (2023) suggests that menopause is actually a widespread trait of mammals, which would change our understanding of senescence and have implications for the study of menopause in humans. Here, we identify three main issues with the methodology of this commentary: the use of captive populations, the use of maximal lifespan, and misinterpretations of the data sources. We show that this methodology does not support the conclusions at the species-level, and conclude that, in line with the predictions of classic life-history theory, menopause is indeed a rare trait.

## INTRODUCTION

In most mammals, reproduction declines gradually with age as a result of the normal ageing process^1,2^. Whilst most females tend to die in nature before reaching complete cessation of reproductive function, in some species, experiencing menopause and having a long post-reproduction lifespan thereafter is common. The existence of a prolonged post-reproductive life has been a long-standing puzzle in evolutionary biology, as classic life history theory predicts that natural selection should favour reproduction until the end of life, in order for organisms to maximise their biological fitness most effectively^3-5^. There has been a lot of confusion in the literature as to the taxonomic prevalence of post-reproductive lifespans as a species-level trait (see ^5^). This debate has been affected by inconsistencies and problems with how post-reproductive lifespans have been measured and by looking at captive populations, which can have artificially long lifespans and artificially early reproductive termination (see ^5^ for a discussion). Thus, to understand how evolution has shaped post-reproductive lifespans, it is essential to look at wild populations where individuals experience natural mortality and fecundity. In 2018 Ellis and colleagues^6,7^ used statistically valid measures on wild populations^8^ to determine that extended post-reproductive lifespans in mammals living under natural conditions are limited to humans and a few species of toothed whale. Following these studies, the debate is considered resolved in the evolutionary literature - extended life after reproduction is a rare taxonomic trait.

In contrast to this resolution, Winkler & Goncalves^9^ put forward the provocative claim that there is a “*false yet widespread belief that cessation of reproductive potential in midlife is limited to humans and a few toothed whales*.” Pooling data from studies on predominantly captive populations, and in contrast to recent evolutionary literature, they claim oopause (their term for permanent age-associated cessation of ovulation) and a post-reproductive life to be widespread among mammals. The results of Winkler & Goncalves^9^ are consistent with previous work, which has shown that under captive conditions, a range of mammalian species in a number of orders do show some degree of post-reproductive life^8,10^. However, Winkler & Goncalves^9^ dismiss the importance of wild data. Given sufficient medical support, protection and food provision, it is not surprising that females in captive populations can live beyond their reproductive years. While this may provide an interesting opportunity to study the physiology of the ageing process, post-reproductive life in captivity fundamentally differs from post-reproductive life in the wild and requires no evolutionary explanation. In contrast, prolonged post-reproductive life seen in wild populations of humans and a handful of mammals challenges classic evolutionary theory and for over half a century, evolutionary biologists have been developing and testing hypotheses to explain why and how post-reproductive life has evolved and why it is such a rare taxonomic trait (see ^5^ for a review). The conclusions and approach for data handling and analysis used by Winkler & Goncalves^9^ risk misinforming those unfamiliar with the evolutionary literature. We feel it is imperative that the state of the field in evolutionary biology is clarified, particularly for those working on menopause from medical and cellular perspectives.

We have identified three key elements of Winkler & Goncalves^9^ study that do not lead us to the same conclusions as the authors, both theoretically and methodologically. We first compare data from Winkler & Goncalves^9^ to that from Ellis et al.^6^ to show that using data from captive populations will not represent wild populations adequately. We then focus our discussion of the comparison of our findings with the findings of Winkler & Goncalves^9^ along three main lines, which are based on core literature in the field: *i*. the use of predominantly captive populations and the exclusion of wild populations, *ii*. the use of an individual-level measure of maximal lifespan for investigating a population-level trait (and more generally the methodology), *iii*. misinterpretations of the literature leading to serious flaws in the data used.

## METHODS

Data used in this study were taken from the supplementary information of Winkler & Goncalves^9^ and from Ellis et al.^6^. To make our work directly comparable, maximum lifespan was defined from life tables used in Ellis et al.^6^ as the age at which survival reached 0. Please note however, we do not recommend this as an approach for comparing species’ life history, and instead we advocate for the use of post-reproductive representation^8^.

## RESULTS

Using information on species from Winkler & Goncalves^9^ and Ellis et al.^6^, we recreated Figure 2 from Winkler & Goncalves^9^ to show how captive populations are not comparable to wild counterparts. There was direct overlapping data for six species (humans, sheep (*Ovies aries*), African elephants (*Loxodonta africana*), Japanese macaques (*Macaca fuscata*), European badgers (*Meles meles*), and chimpanzees (*Pan troglodytes*)), as well as data on similar species in two cases: baboons and gorillas. Wild population data from sheep come from the extensively studied Soay sheep population in the United Kingdom. We use a specific species of baboon (olive baboons *Papio anubis*) rather than *Papio sp*. (which appears to be information on *Papio hamadryas* and *Papio anubis*). Winkler & Goncalves^9^ also used Western gorillas (*Gorilla gorilla*) whereas we use Eastern gorillas (*Gorilla beringei*), though this is unlikely to drive differences in our findings: whilst ecological differences exist between the gorilla species, life history patterns are similar^11,12^. Figure 1a shows that using wild data gives a much lower maximum lifespan post-reproductively for all comparable species. This is due to reduced longevity in wild conditions (Figure 1b), and in some cases shorter reproductive capability in captivity (Figure 1c). In the case of sheep and African elephants, age at oopause (as per Winkler and Gonzales^9^) appears to occur years before the actual age at last birth in wild populations (from ^6^), which logically requires the physiological capability to reproduce. For sheep, the age of the oldest female who gave birth in the wild is 6 years after the age of oopause, and for African elephants the difference is 27 years (Figure 1c).

**Figure 1.**
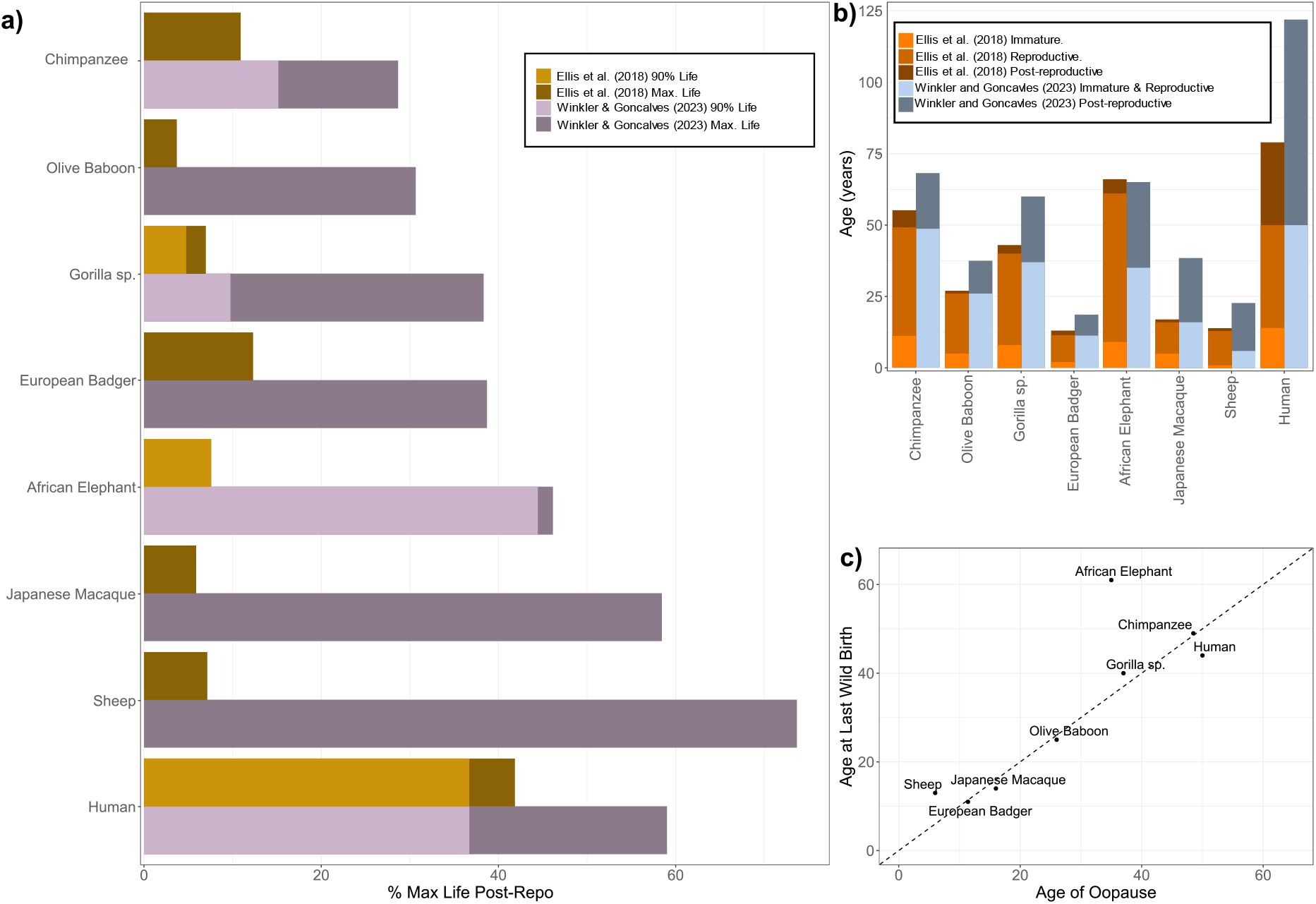
Comparison of wild and captive data. a) For the eight species with comparable data, % Maximum Lifespan post-reproductive is considerably higher in the predominantly captive populations in from Winkler & Goncalves^9^ – purple/grey bars – than from the wild populations in Ellis et al.^6^ - dark yellow bars. Purple/grey bars recreate Figure 2 from Winkler & Goncalves^9^. Bars show the percentage of maximum lifespan (darker bar) or 90% lifespan (lighter bar) after the end of reproduction/’oopause’. For purple/grey bars in some cases 90% lifespan data are not available (Winkler & Goncalves^9^); and for the dark yellow bars 90% lifespan post-reproductive is 0, hence the absence of light yellow bars. b) Comparing the life history of wild (orange bars) and predominantly captive populations (dark blue-grey bars) of eight mammal species. Bar shade represents the life stage of females of a given age, darker colours show later life stages (due to differences in structure of data, no pre-reproductive stage is presented for Winkler & Goncalves^9^). In most species, age of reproductive cessation is comparable for wild and captive populations, but in the predominantly captive data used in Winkler and Goncalves^9^ the post-reproductive period is prolonged compared to the wild data from Ellis et al.^6^. c) Comparison of age at last known birth (from the sources in Ellis et al.^6^) and age at oopause (from Winkler & Goncalves^9^). Species above the dashed diagonal line have an oldest known birth later than age at oopause.

Post-reproductive representation (PrR) is a statistically robust and reliable method^8,10^ for comparing populations that differ in their total lifespans, and is independent of mortality of pre-reproductive individuals^8^. The value indicates how long a female could expect to live post-reproductively in a population e.g. a PrR of 0.3 indicates a female would expect to live an average of 30% of life post-reproductively. As data from Ellis et al.^6^ in Figure 1a uses maximum lifespan from wild populations, the values are not comparable measures to PrR as they do not account for the proportion of years of adult life lived by females post-reproductively in a population. In other words, PrR values would be lower (see Ellis et al.^6^). Of these eight species compared here, only humans have a significant post-reproductive representation (Ellis et al.^6^).

## DISCUSSION

### The use of data on captive vs wild populations

The clear discrepancies between wild and captive populations shown in Figure 1 lead to the first main issue with the conclusions drawn in Winkler & Goncalves^9^: caution is needed when making evolutionary conclusions from captive populations. Whilst studying populations in captivity may be of interest for investigating the physiological basis of reproductive cessation and the maximum theoretical lifespan of species, the evolutionary origin of traits can only be discussed in the context of conditions where they have evolved by natural selection. Without natural selection from extrinsic factors, captive populations are unrepresentative of their species natural populations and do not follow the normal processes of senescence one can observe in the wild^1,13^. As long post-reproductive periods do not occur in most wild populations – the conditions where they evolved – we cannot draw conclusions about how, why and when post-reproductive lifespans evolve from looking at captive populations.

Captivity is a protected environment, where most mammalian species live longer and have a later onset of senescence than in the wild due to removal or reduction of the risk of predation, starvation, disease, and other forms of extrinsic mortality^14^. In addition, reproduction may be negatively affected by captivity for some species^15-20^ which can artificially shorten the reproductive lifespan. Increasing lifespan and shortening reproduction will inevitably lead to significant periods of post-reproductive life being found. Levitis et al.^10^ showed mathematically that protected environments (e.g. captivity) can lead to increased post-reproductive survival, but that “*a species should have significant* [post-reproductive representation, PrR] *across the environments it occupies to conclude that it has a* [post-reproductive lifespan]”; in other words, finding a post-reproductive lifespan in captivity does not mean that the species in question should be classed as having physiological reproductive cessation and a subsequent post-reproductive lifespan^6,10^. Even under the harshest conditions, such as those experienced by plantation slaves in Trinidad, humans show significant post-reproductive lifespans^10^. The conclusions of Winkler & Goncalves^9^ are based on almost exclusively captive populations that cannot be viewed as representative of these species. The authors suggest that captive populations are preferable as “*Populations in which most individuals die substantially earlier than their potential maximum lifespan are particularly problematic, as this will severely underestimate the prevalence of oopause*”, but if individuals in a population rarely reach the age of physiological reproductive cessation, then it should not be considered as a trait of the population and it has no realised evolutionary relevance.

The supplementary of Winkler & Goncalves^9^ lists species, the data used, the data source, and whether that species was included in the paper. Worryingly, very few wild populations were included in this data, even before the exclusion criteria. The tree of life (Figure 1 of Winkler & Goncalves^9^) would not appear to support the argument that physiological reproductive cessation/oopause is a general mammalian trait were the results from wild populations also included (see Ellis et al.^6^ for 50 species with sufficient data that do not show extended post-reproductive life). The only wild populations that were included in Winkler & Goncalves^9^ were four whale species that have previously been shown to have extended post-reproductive life from physiological and demographic data^7,21^, and the European badger^22^.

Despite their unsuitability for evolutionary studies of post-reproductive lifespan, captive populations can have value for investigating whether mechanisms of ageing are generalisable across environments, which could allow the development of environmentally robust ageing interventions^1^. Furthermore, identifying significant post-reproductive representation in unnatural environments (i.e. free from extrinsic mortality pressures) can show that maximum lifespan observed in the wild is not genetically inflexible^8^.

### The use of an individual-level measure of maximal lifespan

Our second main issue with the conclusions in Winkler & Goncalves^9^ is the use of maximum lifespan and the methodology in general. To determine whether a population exhibits a post-reproductive lifespan, one should use population measures that are robust to demographic outliers^8,23,24^ to avoid extreme individual values causing over- or under-estimation of population-level values. Instead, Winkler & Goncalves^9^ predominantly used maximum longevity listed for the species, an individual measure that may be unrepresentative of realised longevity in species^24^. This can be most clearly illustrated with humans, where the maximum lifespan is 122, yet life expectancy is in the mid-80s even in the most developed countries^25^. The methodology employed - comparing an age at reproduction cessation (but see below for issues with some of this data) to the maximum known longevity or when 90% of the animals in the population are deceased - is not a validated method, and appears to be a simplification of post-reproductive time (the proportion of life lived post-reproductively). The deficiency in this methodology is evident: assume that species A reproduces until death (i.e. no reproductive cessation), with the vast majority of the population dying by age 20 and one exceptional individual living to an extreme of 30 years. This would result in a supposed 33% of life spent post-reproductively for species A. Hence, the measures used by Winkler & Goncalves^9^ are well-known to be unsuited for demonstrating the evolutionary relevance of post-reproductive life^8^.

Additionally, Winkler & Goncalves^9^ suggest that any physiological measures are better indicators of the end of reproductive life than demographics, but this is not true on a population level. If the end of reproduction calculated from demographic information does not lead to a finding of a significant post-reproductive period, then using physiological markers of reproductive cessation will not make a difference as they will either be at the age of reproductive cessation on a population-level or after it. Physiological measures indicating end of reproductive period in most individuals before the population-level age at last reproduction (such as those supposedly found for African elephants in Figure 1; see section on misinterpretations in the literature below for details) raises the question of whether such markers are relevant for the species, as continued reproduction after supposed physiological cessation is a clear indicator that physiological cessation has not occurred. The only cases where knowing about the physiological cessation of reproduction would be useful in this context are for cases such as the Asian elephant (*Elephas* maximus) to determine whether the significant PrR^26,27^ is from physiological cessation of reproduction or due to social or other factors that influence mate choice and mating success^27^. PrR should be the preferred methodology, because it is the only one that can distinguish meaningful post-reproductive lifespan from that which occurs by chance. The difference between the two is of evolutionary importance: a short post-reproductive period can occur due to individual variation in the rates of reproductive and somatic senescence^5^, whilst a long, significant post-reproductive period is indicative of a product of selection on that life history.

### Misinterpretations of the literature leading to flaws in the data used

In preparing this response and reviewing the data used by Winkler and Goncalves^9^ we encountered multiple instances where information from published studies was incorrectly used or conclusions were drawn that are not supported by the original study. Firstly, and of large concern, in several cases the age at oopause for species given in Winkler and Goncalves^9^ was considerably younger than the age of the oldest reproducing females (and therefore presumably still producing ova) in the studies the authors cite as sources. For ewes (sheep), for example, the authors use age 6 as the upper age of onset of acyclicity, though in the source paper this can be seen to be the age at which reproduction begins to slow down (in terms of number of lambs being birthed, rather than any physiological measure), and that ewes in the sample continued to reproduce until age 11^28^. To use the same metric in humans would give an age of oopause in the early 30s^29^, well before the actual age at which menopause typically occurs. In combination with the use of maximum age, this can greatly inflate the calculated proportion of life lived post-reproductively. For African elephants, the maximum age at onset of acyclicity is cited to be 35, but the source for this shows that, whilst an absence of reproductive cycling (acyclicity) was highest for 31-35 year old African elephants (47%), the percentage of acyclic individuals actually decreased in the subsequent age groups^30^; in other words, the majority of older females continued to cycle and reproduce. Secondly, in two cases Winkler & Goncalves^9^ directly contradict detailed analysis in the sources they cite. The use of false killer whales (*Pseudorca crassidens*) from Ellis et al.^7^ as supporting the claim of having oopause via ovarian indicators is incorrect: the paper explicitly states that false killer whales were found not to have an extended post-reproductive lifespan on the basis of ovarian markers - although it should be noted that other studies have found convincing support for physiological cessation of reproductive activity before the end of life in female false killer whales^21^. Similarly, the source for wild European badgers investigated the post-reproductive representation of the species, and found that it was insignificant^22^ i.e. an extended post-reproductive lifespan is not a general trait of the species, even if some individuals were shown to have experienced physiological reproductive cessation. In both cases it appears that if physiological cessation is observed for any individual then this has been considered by Winkler & Goncalves^9^ as sufficient evidence of physiological cessation and post-reproductive lifespan for the entire species. The existence of post-reproductive life at an individual level can be due to variation in rates of senescence between individuals, and does not require further evolutionary explanation. Similarly, Asian elephant age at onset of acyclicity is claimed to be 55 years old, despite the original paper and subsequent work on the same population (also cited in the supplementary^9^) proposing that the extended post-reproductive lifespan found was not due to physiological reproductive cessation as many females were still able to breed years after this age, but may instead be a ‘social menopause’^26,27^; in both papers, continued reproduction in older females was shown.

## Conclusion

We agree with the general conclusion of Winkler and Goncalves^9^ that under captive conditions, females will often live to be post-reproductive – we note, however, that the issues we identify above with how the data was included and analysed have likely led to an overestimation of the duration and prevalence of this effect. A key question that needs to be asked is how generalisable these findings on captive populations are to other populations of that species – including those living under natural conditions. When we compare the data of Winkler and Goncalves^9^ to data from wild populations, we find no support that post-reproductive life is a general species trait across a wide range of mammals. Indeed, when using appropriate data analysis methods and including data on wild populations, post-reproductive life as a species-level trait is restricted to humans and a handful of mammals^6,7,21^.

Winkler and Goncalves^9^ ta e issue with using the wider use of the term ‘menopause’ in the literature, (correctly) defining it etymologically as the cessation of menses. However, as researchers working on post-reproductive life from an evolutionary perspective, we do not believe that the use of ‘menopause’ is problematic or has caused confusion. Focusing on the physiological process and etymology limits the possible use of the term to humans and the limited number of species exhibiting menstruation (most mammals reabsorb the uterine lining^31^). As highlighted by Winkler & Goncalves^9^, this is unhelpfully limiting and prevents productive and correct interspecies comparisons. However, when used in the evolutionary literature, it more-or-less follows the broader redefinition from Cohen^32^ for cross-species comparative purposes: “*the irreversible loss of the physiological capacity to produce offspring due to intrinsic biological factors*”. ntroduction of novel terminology i.e. ‘oopause’ is, in this case, unnecessary.

Understanding the taxonomic prevalence of long periods of post-reproductive life (relative to lifespan) is vital so that we can identify other species that could be studied to help understand how extended post-reproductive periods could have evolved and been maintained. Long-term individual-based datasets on wild mammal populations, such as killer whales (*Orcinus orca*) that have comparable post-reproductive life to humans, has allowed us to test the generality of the evolutionary framework proposed to explain the evolution of menopause in humans and provides a very rare and informative comparison for human life history evolution^33-39^. While observing that females of the captive population can have artificially long lifespans and post-reproductive life may open up opportunities for new work to study the physiology of the ageing process, captive populations can tell us little about how menopause has evolved. For this, we need to turn our attention to the handful of wild species where females show prolonged life after reproduction.

## AUTHOR CONTRIBUTIONS

Conceptualization, S.N.C., S.E., M.L., D.P.C., and V.L.; Writing – Original Draft, S.N.C.; Writing – Review & Editing, S.N.C., S.E., M.L., D.P.C., and V.L.; Visualization, S.E.

## ACKNOWLEDGMENTS

This work was funded by the Academy of Finland (decision numbers: 320162, 345185, 345183, 331816) and NERC (NE S010327/1).

